# Enhanced analgesic cholinergic tone after neuropathy

**DOI:** 10.1101/2020.06.11.146852

**Authors:** Dhanasak Dhanasobhon, Maria-Carmen Medrano, Yunuen Moreno-Lopez, Sehrazat Kavraal, Charlotte Bichara, Rémy Schlichter, Perrine Inquimbert, Ipek Yalcin, Matilde Cordero-Erausquin

**Affiliations:** Institut des Neurosciences Cellulaires et Intégratives, Centre National de la Recherche Scientifique, 67000 Strasbourg, France; Université de Strasbourg, Strasbourg, France; University of Strasbourg Institute for Advanced Study (USIAS), Strasbourg, France

**Author notes:** Corresponding author: Matilde Cordero-Erausquin, PhD, HDR, CR1 CNRS, Institut des Neurosciences Cellulaires et Intégratives - CNRS UPR 3212, 8 allée du Général Rouvillois, 67000 Strasbourg cedex, France.

**Keywords:** behavior, cholinergic system, dorsal horn, electrophysiology, mechanical stimuli, mice, neuropathic pain, nicotinic receptors, spinal cord, von-Frey test

## Abstract

At the spinal cord level, a tone of endogenous acetylcholine (ACh) modulates nociceptive sensory processing. Increasing the level of spinal ACh induces analgesia in naïve animals or in situation of acute pain in the clinics, but whether this is still the case in situations or models of chronic pain is controversial. Here, we demonstrate the persistence, and even increased impact of the analgesic cholinergic tone acting through nicotinic receptors in neuropathic mice. The neuropathy does not affect the number and properties of dorsal horn cholinergic neurons, proposed to be the source of spinal ACh. Subthreshold doses of acetylcholinesterase (AChE) inhibitors in sham animals become anti-allodynic in cuff mice suggesting that the alterations occur in the cholino-receptive neurons. Thus endogenous cholinergic signaling can be manipulated with low doses of drugs to relieve mechanical allodynia in animal neuropathy models. This opens new avenues with potentially fewer side effects for neuropathic pain treatment.

## 1. Introduction

Endogenous acetylcholine (ACh) is an important modulator of nociceptive sensory processing in the spinal cord. Epidural administration of neostigmine, an acetylcholinesterase (AChE) inhibitor that induces an increase in ACh spinal level by preventing its degradation, produces pain relief for child birth and post-operation pain in clinics (Eisenach, 2009). In rodents, local spinal injection of AChE inhibitors (by intrathecal injections, or i.t.) similarly produces analgesia to thermal or chemical noxious stimulation (Chen & Pan, 2003; Chiari, Tobin, Pan, Hood, & Eisenach, 1999; Hartvig, Gillberg, Gordh, & Post, 1989; Miranda, Sierralta, & Pinardi, 2002; Naguib & Yaksh, 1994). The analgesic effect of endogenous ACh is not only observed when its level is artificially increased (by AChE antagonists), but also with its physiological levels. Indeed, impairment of cholinergic signaling by locally antagonizing ACh receptors (nicotinic or muscarinic) or by knocking-down the β2 subunit of nAChRs induces hyperalgesia and/or allodynia in rodents (Hama & Menzaghi, 2001; Rashid & Ueda, 2002; Yalcin, Charlet, et al., 2011; Zhuo & Gebhart, 1991). This suggests that a basal “tone” of spinal ACh modulates the nociceptive threshold.

A dense plexus of cholinergic processes (composed of both dendrites and axons) lies in laminae (L) II-III of the dorsal horn (DH) of rodents (Barber, et al., 1984; Mesnage, et al., 2011). Within this plexus, cholinergic dendrites receive synaptic contacts from primary afferents, and cholinergic axons make presynaptic contacts onto them (Olave, Puri, Kerr, & Maxwell, 2002; Ribeiro-da-Silva, 2004), offering a potential substrate for sensory modulation. In rodents, sparse cholinergic neurons with extended ramification, are located in lamina III-IV and have been proposed to be the source of this plexus. They could therefore be responsible for cholinergic analgesia (Mesnage, et al., 2011). Interestingly, we have demonstrated that a similar cholinergic population exists in the dorsal horn of macaque monkeys, with comparable localization, density and most importantly, synaptology (Pawlowski, et al., 2013). The behavioral and anatomical similarities have confirmed that mice are a valuable model to study cholinergic analgesia. But how this population of cholinergic interneurons ultimately impacts nociceptive behavior is still unknown.

AChE inhibitors are used in the clinics only in a context of acute pain. Yet chronic pain is also a devastating and widespread problem, inadequately treated according to two-thirds of chronic pain sufferers (Breivik, Collett, Ventafridda, Cohen, & Gallacher, 2006). How the endogenous spinal cholinergic control of analgesia evolves in situations of chronic pain in primate is unknown, and data on rodents are scarce and at times contradictory. While Rashid and Ueda have proposed that the cholinergic tone is disrupted in neuropathic animals as shown by the loss of effect of the i.t. injection of a nicotinic antagonist (Rashid & Ueda, 2002), i.t. injections of AChE antagonists still have an analgesic effect (Hwang, et al., 1999; P. Lavand’homme, Pan, & Eisenach, 1998; P. M. Lavand’homme & Eisenach, 1999; Takasu, Honda, Ono, & Tanabe, 2006) suggesting that ACh is still locally available to affect sensory processing. The dorsal horn of the spinal cord is a site for numerous forms of plasticity occurring during chronic pain (Tsuda, Koga, Chen, & Zhuo, 2017), and it was recently demonstrated that synaptic inputs or intrinsic properties of several interneuronal populations are altered during the neuropathy (Cheng, et al., 2017; Imlach, Bhola, Mohammadi, & Christie, 2016; Peirs, et al., 2015; Petitjean, et al., 2015). How the cholinergic population, and more generally the spinal cholinergic analgesia, is affected in chronic pain models remained to be evaluated and was the main goal of this study. Accordingly, we have combined behavioral, histological, *in vivo* and *in vitro* electrophysiological recordings to improve our understanding of spinal cholinergic analgesia. We specifically addressed the question of its plasticity in conditions of chronic pain, and of potential underlying mechanisms.

## 2. Results

### 2.1 Behavioral effect of cholinergic modulation onto mechanical responses

We first characterized the dose-dependent effect of spinal cholinergic modulation onto transmission of mechanical information in naïve CD1 mice. For this purpose, the mechanical paw withdrawal threshold (PWT) was evaluated with the von Frey test, before and after i.t. injection of the AChE inhibitor physostigmine. Physostigmine (15 nmol, i.t), produced a long-lasting increase in the mechanical withdrawal threshold (Fig. 1A, Suppl. Fig. 1; statistics are presented in the figure legends) while lower doses (1.5-7.5 nmol) were without effect. This indicates that ACh is endogenously released and its level in the spinal cord closely controlled by AChE. We thus confirm that, if its concentration is increased (by inhibiting its hydrolysis), ACh reaches downstream targets, activating a spinal circuit leading to analgesia.

**Figure 1:**
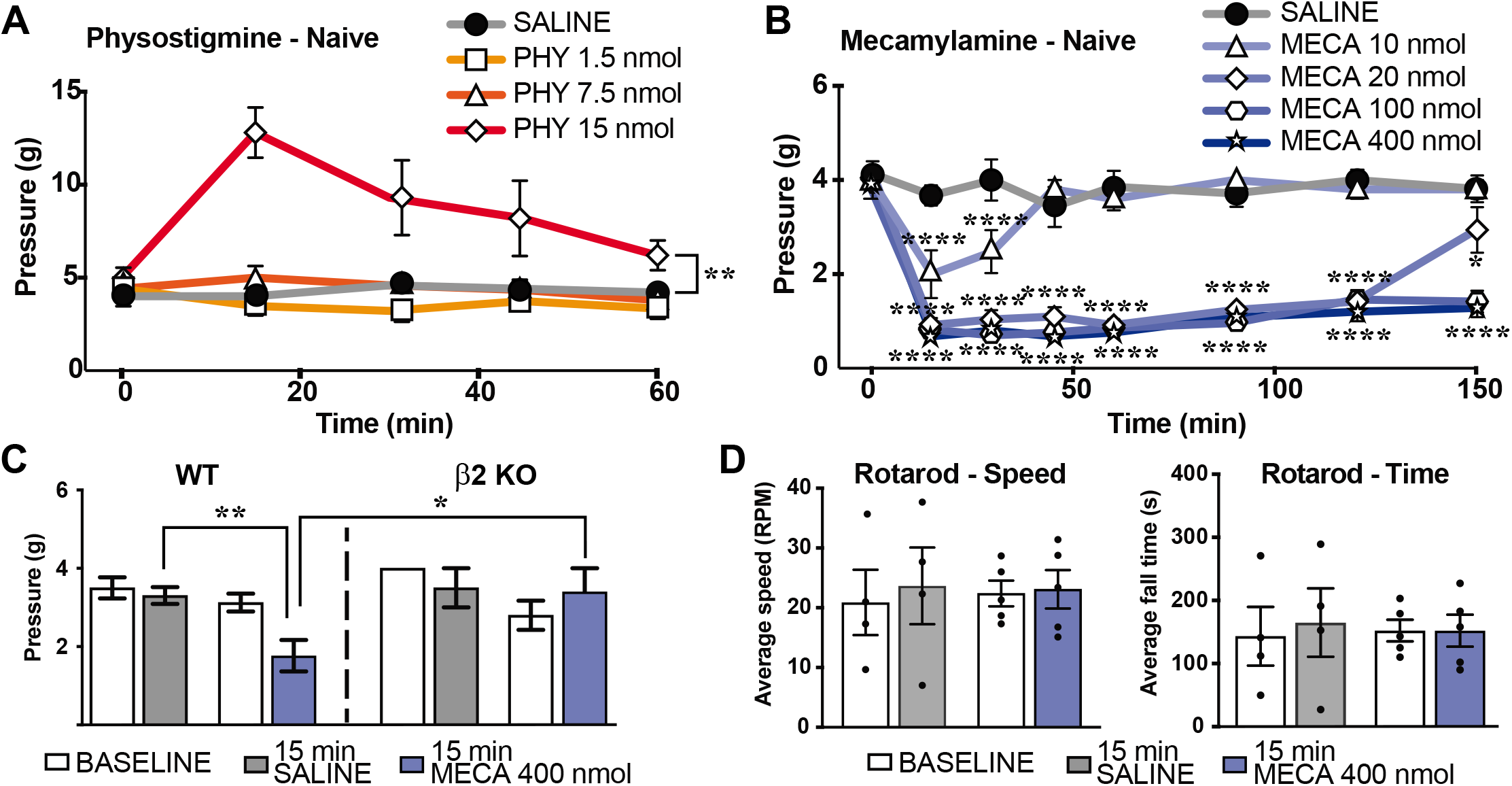
Time course of the effect of cholinergic drugs on mechanical threshold and motor coordination in naive animals. A: Effect of i.t. injection of an AChE inhibitor, physostigmine, on the paw withdrawal threshold in the Von Frey test. 15 nmol physostigmine produced an analgesic effect in naïve mice (Groups of n = 5 mice, Drug effect: p=2.43e-3, nparLD). ** p < 0.01 mean of PHY 15 nmol vs. mean of saline, Bonferroni’s multiple comparisons test. B: Effet of i.t. injection of a nicotinic antagonist, mecamylamine, on the paw withdrawal threshold in the Von Frey test. Mecamylamine induced mechanical allodynia at 10, 20, 100 and 400 nmol in naive CD1 mice (Groups of n = 5-7 mice, Drug-Time interaction: p = 3.30e-12). **** p < 0.0001, *** p < 0.001, ** p < 0.01, * p < 0.05, vs. saline, Bonferroni’s multiple comparisons test. In 1B, the bottom asterisks concern the two higher doses. C: Effet of i.t. injection of mecamylamine, on the paw withdrawal threshold in the Von Frey test in WT and β2* nAChR knock-out (β2 KO) C57BL/6 mice (Groups n = 2 – 10 mice, Strain-Drug-Time interaction: p = 0.022, nparLD). Baseline is the PWT before any injection, and the PWT measured 15 minutes after the injection is presented for the saline and and 400 nmol mecamylamine group. Mecamylamine induced mechanical allodynia at 400 nmol in WT but not β2 KO mice (Strain effect: p = 0.028, Drug effect: p = 0.001, Time effect: p = 6.25e-4, Strain-Drug-time interaction: p = 0.022, nparLD). D: Effet of i.t. injection of 400 nmol mecamylamine on rotarod performances. Mecamylamine did not change speed of fall or time of fall in naïve CD1 mice (Groups n = 4 mice, Speed - Drug effect: p = 0.925, Time effect: p = 0.506, Drug-Time interaction: p = 0.541; nparLD | Time - Drug effect: NA, Time effect: p = 0.535, Drug-Time interaction: p = 0.467; nparLD).

In order to investigate whether this cholinergic analgesic mechanism is active in baseline physiological conditions, we interfered with it by blocking cholinergic nicotinic receptors. We thus injected the non-specific nicotinic antagonist mecamylamine (10-400 nmol, i.t.) and tested at different time points after the injection (Fig. 1B). Mecamylamine induced a dose- and time-dependent reduction in the PWT: for all doses, the PWT was statistically reduced as soon as 15 min post-injection, and for all but the 10 nmol dose, the effect lasted for more than two hours. In the 15 min – 2 hours time window, there was no significant difference between the three higher doses (at each time point, 400 nmol or 100 nmol vs. 20 nmol: p>0.99, Bonferroni’s multiple comparisons test) suggesting that the effect already plateaued with 20 nmol mecamylamine. 150 minutes after the injection, however, animal that had received 20 nmol of mecamylamine had a PWT statistically different from the one of animals that had received 100 or 400 nmol mecamylamine (20 nmol vs. 100 nmol, p=0.0007; 20 nmol vs. 400 nmol, p=0.0001, Bonferroni’s multiple comparisons test). Indeed, the allodynia evoked by 10 nmol mecamylamine had faded away after 150 minutes, while it was still present in the 20, 100 and 400 nmol injection groups (comparisons with saline presented in Fig. 1B). This suggests that the cholinergic analgesic mechanism is active in baseline physiological conditions, either because ACh is continuously released and activating downstream circuits, or because it is released as a consequence of the mechanical stimulation. Whatever the underlying cause, the pharmacological effect of a nicotinic antagonist defines the presence of an analgesic spinal cholinergic tone, acting via nicotinic receptors.

High concentrations of mecamylamine are known to have potentially non-specific effects (McDonough & Shih, 1995; O’Dell & Christensen, 1988). We used β2 nicotinic knock-out mice (β2 KO) to verify whether the highest dose of mecamylamine used (400 nmol) was acting specifically through inhibition of nAChR receptors. While 400 nmol i.t. mecamylamine induced a marked reduction of the PWT in WT animals, it produced no effect on β2 KO mice (Fig. 1C, 400 nmol mecamylamine vs saline in WT mice: p = 0.0042, and in KO: p>0.99, Bonferroni’s multiple comparisons test). Moreover, we observed that this high concentration of mecamylamine did not affect motor-coordination as evaluated by the rotarod test in naive mice (Fig. 1D), suggesting that the observed modulation strictly involved mechanical sensory processing.

### 2.2 The spinal cholinergic tone is present after neuropathy

We then investigated the alteration of this analgesic cholinergic tone in a mouse model of neuropathy. We used the cuff model and analyzed the effect of i.t. injection of mecamylamine (10-400 nmol) on the PWT of sham and cuff mice. For the ipsilateral, but not the contralateral, paw there was a significant interaction between Surgery, Time and Drug (p = 0.0354), indicating that the neuropathy modified the response of the ipsilateral paw to mecamylamine.

Mecamylamine induced a dose-dependent reduction in the mechanical withdrawal threshold of the contralateral hindpaw in sham and cuff mice (Fig. 2A Top, in sham, for all doses of mecamylamine vs. saline at 15 min, p<0.0001 in post-hoc tests; in cuff, similar comparison yielded to p<0.0001 for the 20 nmol, 100 nmol and 400 nmol mecamylamine doses, and p = 0.0257 for the 10 nmol dose, Bonferroni’s multiple comparisons test). For the ipsilateral hindpaw, 10 and 20 nmol was ineffective in cuff as opposed to sham groups (Fig. 2A Bottom, post-hoc tests at 15 min, drug vs. saline: p = <0.0001 for 10 and 20 nmol mecamylamine in sham and respectively p>0.99 and p=0.159 in cuff; Bonferroni’s multiple comparisons test). The mechanical allodynia observed in cuff animals was exacerbated when higher doses were injected (Fig. 2A; post-hoc tests at 15 min, in cuff mice: for 100 nmol mecamylamine vs. saline, p = 0,0017 and for 400 nmol mecamylamine vs. saline, p<0.0001; Bonferroni’s multiple comparisons test). Analysis of the dose-response curve demonstrates a shift to the right in cuff mice, with higher doses being necessary to obtain a significant effect (Fig. 2B; Ipsilateral side - Least square fit, p<0.001). Thus the spinal cholinergic tone is still present in cuff animals, involving nAChRs and affecting mechanical responses. The shift to the right of the mecamylamine dose-response curves further suggests that the cholinergic tone might be increased by the pathology, either because more ACh is released or because endogenous ACh has an increased effect on its post-synaptic targets. We next investigated these two possibilities.

**Figure 2:**
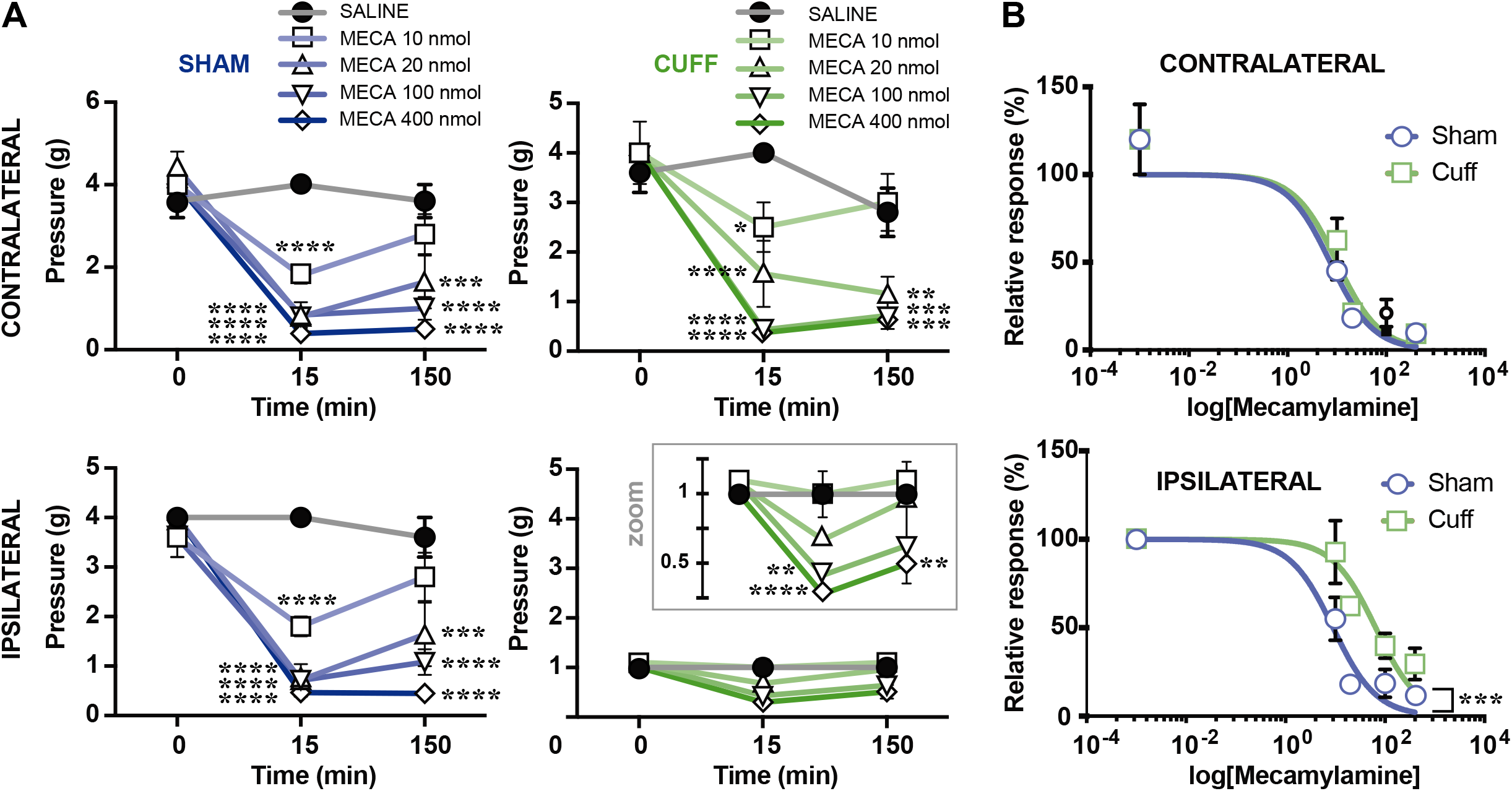
Mecamylamine shows that a spinal cholinergic tone is still present after neuropathy. A: Time course of the effect of intrathecal mecamylamine on mechanical threshold of sham and cuff mice (groups n=4-5). For the contrateral side (Top panels), there was a significant effect of the Drug (saline, 10-400 nM mecamylamine, p = 1.901e-19, nparLD) and Time (0, 15 and 150 min post i.t. injection, p = 1.95e-67, nparLD) but not of the Surgery (sham vs. cuff, p = 0.702, nparLD), as well as a significant interaction between Drug and Time (p = 1.79e-16, nparLD). For the ipsilateral side (Bottom panels), there was a significant effect of Surgery (p = 8.64e-19), Drug (p = 6.36e-15) and Time (p = 3.07e-53), as well as a significant interaction between Drug and Time (p = 1.94e-16) and between Surgery, Time and Drug (p = 0.0354). While treatment with 10 and 20 nmol mecamylamine had no effect, 100 and 400 nmol mecamylamine further potentiated the mechanical allodynia in the ipsilateral paw. **** p < 0.0001, *** p < 0.001 ** p < 0.01, * p < 0.05 vs. saline, Bonferroni’s multiple comparisons test. B: Dose response curve of mecamylamine on responses of sham and cuff animals. Neuropathy induced a shift to the right to the response to mecamylamine (Ipsilateral side: sham vs. cuff p<0.001; Least square fit).

### 2.3 Plasticity of LIII/IV cholinergic interneurons during neuropathy

Lamina III-IV cholinergic neurons are the most plausible source of the spinal cholinergic tone. We thus closely analyzed this population in sham and neuropathic mice to test the hypothesis that the release of ACh could be increased following the neuropathy. In particular, we assessed 3 different criteria: (1) synaptic inputs, (2) active and passive electrophysiological properties, and (3) number of cholinergic interneurons.

#### 2.3.1 Synaptic inputs to lamina II and lamina III-IV interneurons

We then assessed whether DH cholinergic neurons could be differentially activated in sham vs. cuff by comparing the synaptic inputs received by these neurons in these two conditions. Cholinergic interneurons are located in lamina III-IV, and best preserved in horizontal sections (Mesnage, et al., 2011). However, we first performed a set of recordings of lamina II neurons in transverse slices, in order to put our results in context with the literature, mostly focusing on this layer. Indeed, while the cuff model of mouse peripheral neuropathy has been well described in behavioral terms (Benbouzid, et al., 2008; Yalcin, Bohren, et al., 2011), we were the first to analyze the plasticity occurring in the spinal cord with electrophysiology *in vivo* (Medrano, Dhanasobhon, Yalcin, Schlichter, & Cordero-Erausquin, 2016) and now *in vitro*. Recordings were performed in the ipsilateral (to cuff insertion) DH in the presence of 0.5 μM TTX (Fig. 3A). The frequency of miniature IPSCs was lower in cuff mice compared to sham controls, whereas the frequency of miniature EPSCs was unchanged (Fig. 3B). The amplitude for miniature currents (either EPSC or IPSCs) were similar for sham and cuff mice (Fig. 3B). Thus only the frequency of mIPSCs is modified in lamina II neurons in cuff animals, similarly to what is observed in the spared nerve or chronic constriction injury models (Iura, Takahashi, Hakata, Mashimo, & Fujino, 2016; Moore, et al., 2002).

**Figure 3:**
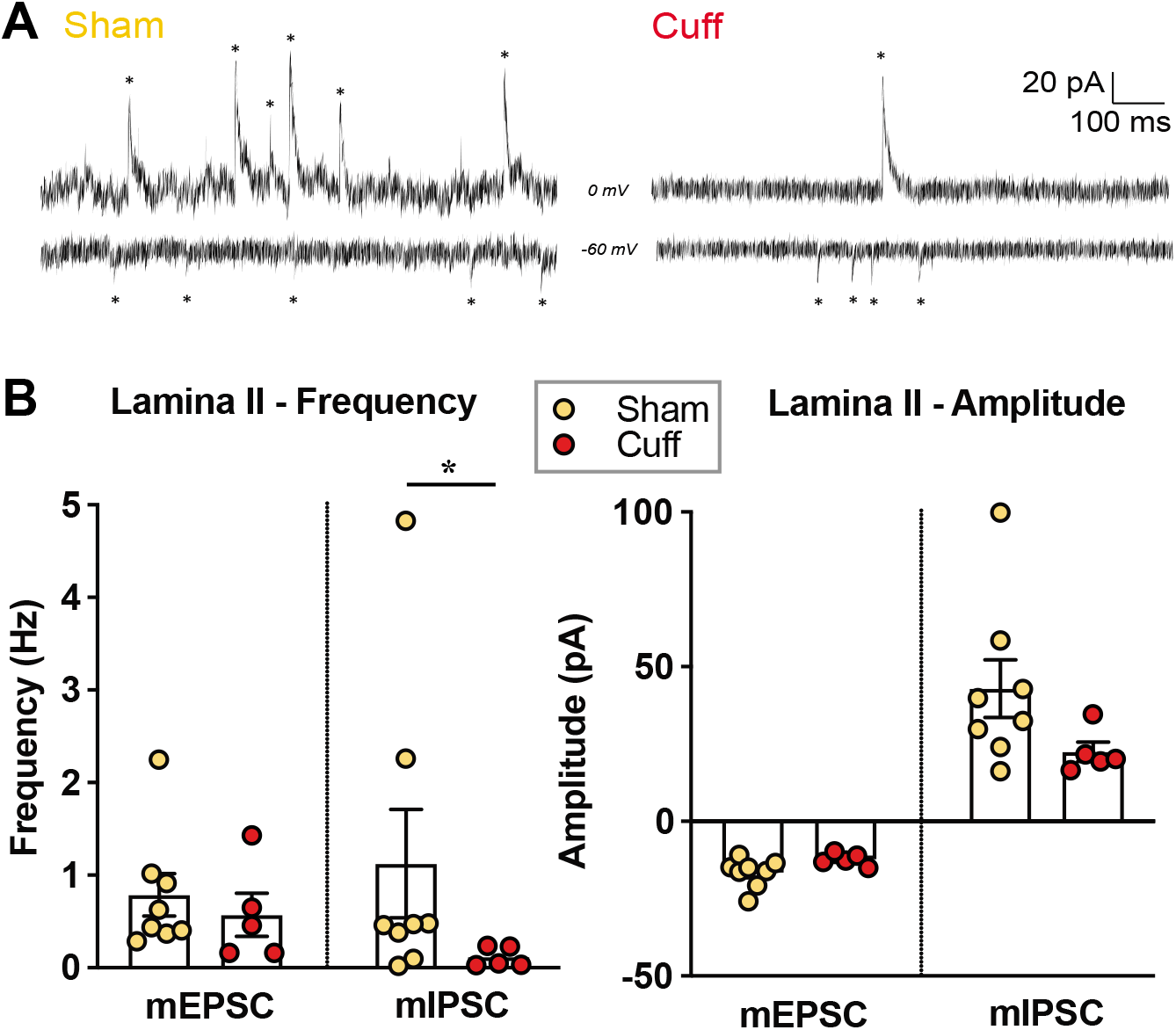
In vitro electrophysiological characterization of the cuff model. A: Representative patch-clamp recordings from LII neurons illustrating miniature excitatory (mEPSC, at −60 mV) and inhibitory (mIPSC, at 0 mV) postsynaptic currents B: (left) Frequency changes in mEPSC and mIPSC recorded LII neurons (n = 5-9 neurons per group): a reduction was observed in mIPSCs frequency (Right) but not mEPSCs (Left) in cuff mice (respectively, p = 0.042 and p = 0.586; Kolmogorov-Smirnov test). (right) Analysis of the amplitude of the same currents demonstrates no effect of the surgery (mEPSC: p= 0.0793; mIPSC: p = 0.0793; Kolmogorov-Smirnov test)

We then moved to lamina III-IV and to horizontal slices to record spontaneous and miniature EPSCs and IPSCs from ChAT::EGFP and non-EGFP (unidentified) neurons in their close vicinity in sham and cuff mice (Fig. 4A, Suppl. Fig. 2). For EPSCs, there were no significant difference between the groups. For IPSCs, there was on average more inhibition in cuff vs. sham mice (not specific to any neuronal population under study, or to any type of current, spontaneous vs. miniatures), and cholinergic neurons had on average less inhibitory inputs than non-cholinergic ones (whether in sham or in cuff mice). There was therefore no change of inputs frequency that was specific to cholinergic neurons after neuropathy (Fig. 4B). We compared with a similar approach the amplitude of miniature EPSC and IPSC in the different neurons and animals, and observed no significant difference between groups (Fig. 4C).

**Figure 4:**
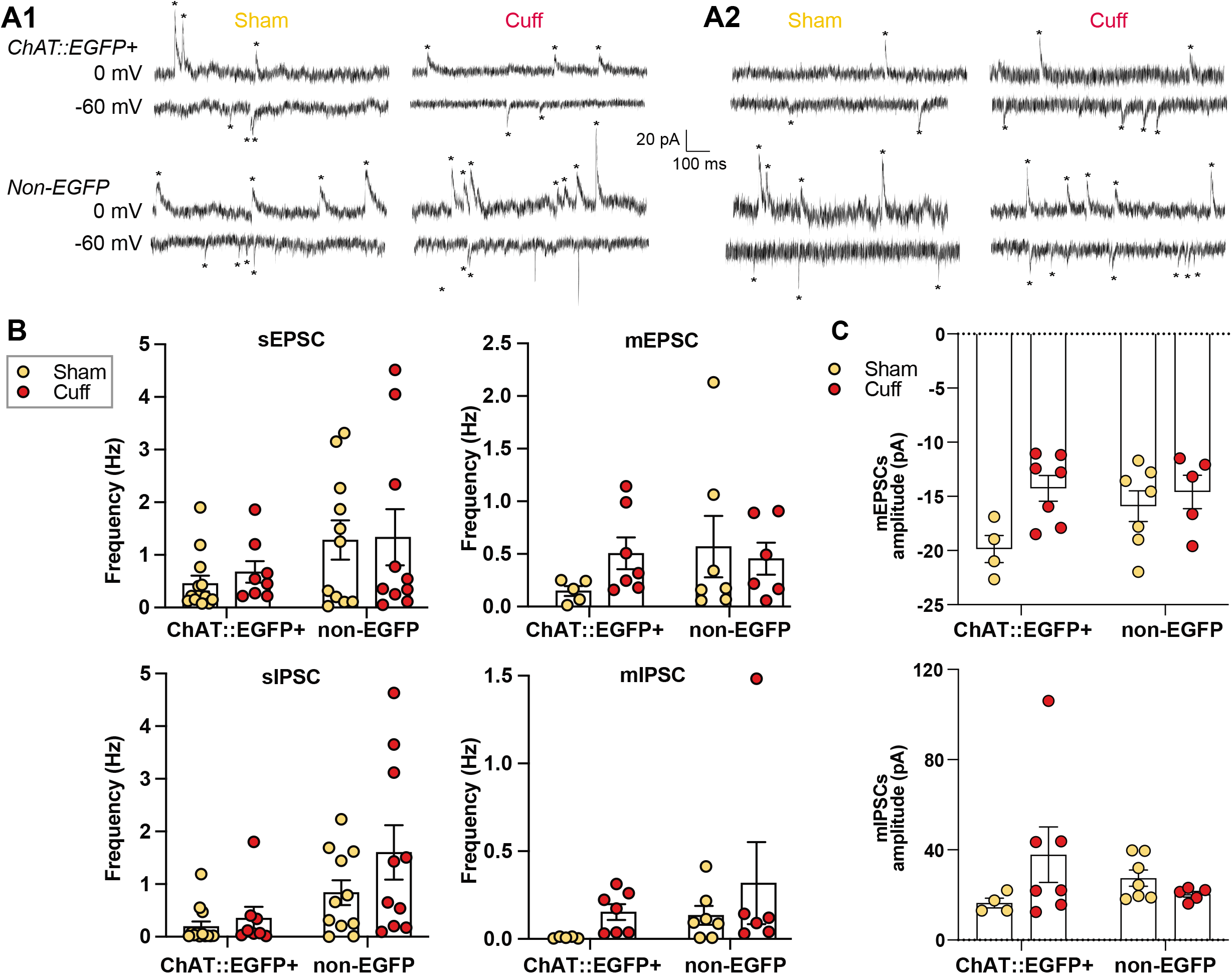
Number and synaptic inputs to Dorsal Horn cholinergic interneurons in sham and cuff mice. A: Representative in vitro patch-clamp recordings from LIII/IV ChAT::EGFP (top) and non-EGFP (bottom) neurons, in Sham and Cuff mice. Recordings are performed at 0 mV to record IPSCs and −60 mV to observe EPSCs. A1: spontaneous excitatory (sEPSC) and inhibitory (sIPSC) postsynaptic currents, A2: miniature mEPSC and mIPSC currents. B: Frequency of spontaneous (left) and miniature (right) EPSCs (top) and IPSCs (bottom) recorded in ChAT::EGFP and non-EGFP cells in horizontal slices of sham and cuff mice (n = 5-13 neurons per group). The distribution of frequencies being non-normal, their log10 was taken to perform the ANOVA. There was no statistical difference in EPSCs frequencies between groups (ChAT::EGFP+ vs. non-EGFP, Sham vs. Cuff, spontaneous vs. miniature). For IPSCs, main effects were significant: neuron (ChAT::EGFP vs. non-EGFP, p = 0.000179, 2-way ANOVA), surgery (sham vs. cuff, p = 0.004616, 2-way ANOVA) and type of current (spontaneous vs. miniatures, p = 1.051e-3, 2-way ANOVA). C: Amplitude of miniature EPSC and IPSC of ChAT::EGFP+ and Non-EGFP+ neurons in sham and cuff mice (n = 4-7 neurons per group). Data was normalized through inversion. There was no signicant difference between groups for mEPSCs (ANOVA). There was a significant Surgery-Neuron interaction for mIPSCs amplitude (p = 0.0437) although this was not confirmed by post-hoc analysis.

These recordings reveal an increase in the frequency of inhibitory inputs that is not specific to lamina III-IV cholinergic neurons as it is also observed in non-EGFP neurons in the same laminae, while lamina II neurons experience a decrease of the frequency of IPSCs. The increased frequency of inhibitory currents would argue, if anything, that cholinergic neurons are less prone to firing (and releasing ACh) in neuropathic conditions. In order to test this hypothesis, we next compared the capacity of cholinergic neurons to be excited by direct depolarization in sham vs. cuff mice.

#### 2.3.2 Active and passive electrophysiological properties

We analyzed the firing patterns of recorded neurons after injection of depolarizing currents, and classified their discharge profiles as either “tonic”, “single-spike”, or “phasic” (cf. Methods, Fig. 5A1).There was no statistical difference between ChAT::EGFP and non-EGFP neurons concerning the proportion of the different firing patterns in the two different animal groups (Fig. 5B). A fraction of recorded neurons presented spontaneous ongoing firing at rest (rheobase of 0pA, Fig. 5C) but their proportion was not different between types of neurons (ChAT::EGFP vs. non-EGFP) or type of surgery (sham vs. cuff) (3×3×3 contingency table, p = 0.2674). Specifically, 2 out of 6 ChAT::EGFP neurons (33.3%) were spontaneously active in sham mice, but none out of 8 (0%) in cuff, while 1 out of 7 non-ChAT neurons (14.2%) was spontaneously active in sham mice vs. 2 out 6 (33.3%) in cuff mice. There also was no statistical difference in the rheobase in the different groups (Fig. 5C). We compared the input-output properties of the different neurons (measured as mean instantaneous firing frequency induced by an increasing depolarizing current, Fig. 5D). On average, there was no effect of the surgery on the instantaneous firing frequencies, but there was an effect of the intensity of the injected current, as well as of the type of neurons: ChAT::EGFP vs. non-EGFP neurons (Suppl. Fig. 3). Moreover, there was an interaction between the response to different injected currents and the type of neurons, as non-EGFP neurons fired more than ChAT::EGFP neurons when large currents were injected (120 to 180 pA, suppl. Fig. 3). There was no interaction of these factors with the surgery, suggesting that the input-output properties of the neurons were not affected by the surgery. We also investigated the passive membrane properties of recorded neurons. There was no difference among groups (ChAT::EGFP or non-EGFP neurons, sham or cuff) in the resting potential (Fig. 5C), the input resistance (Fig 5C) or the amplitude of a sag current (Fig. 5A2, 5E).

**Figure 5:**
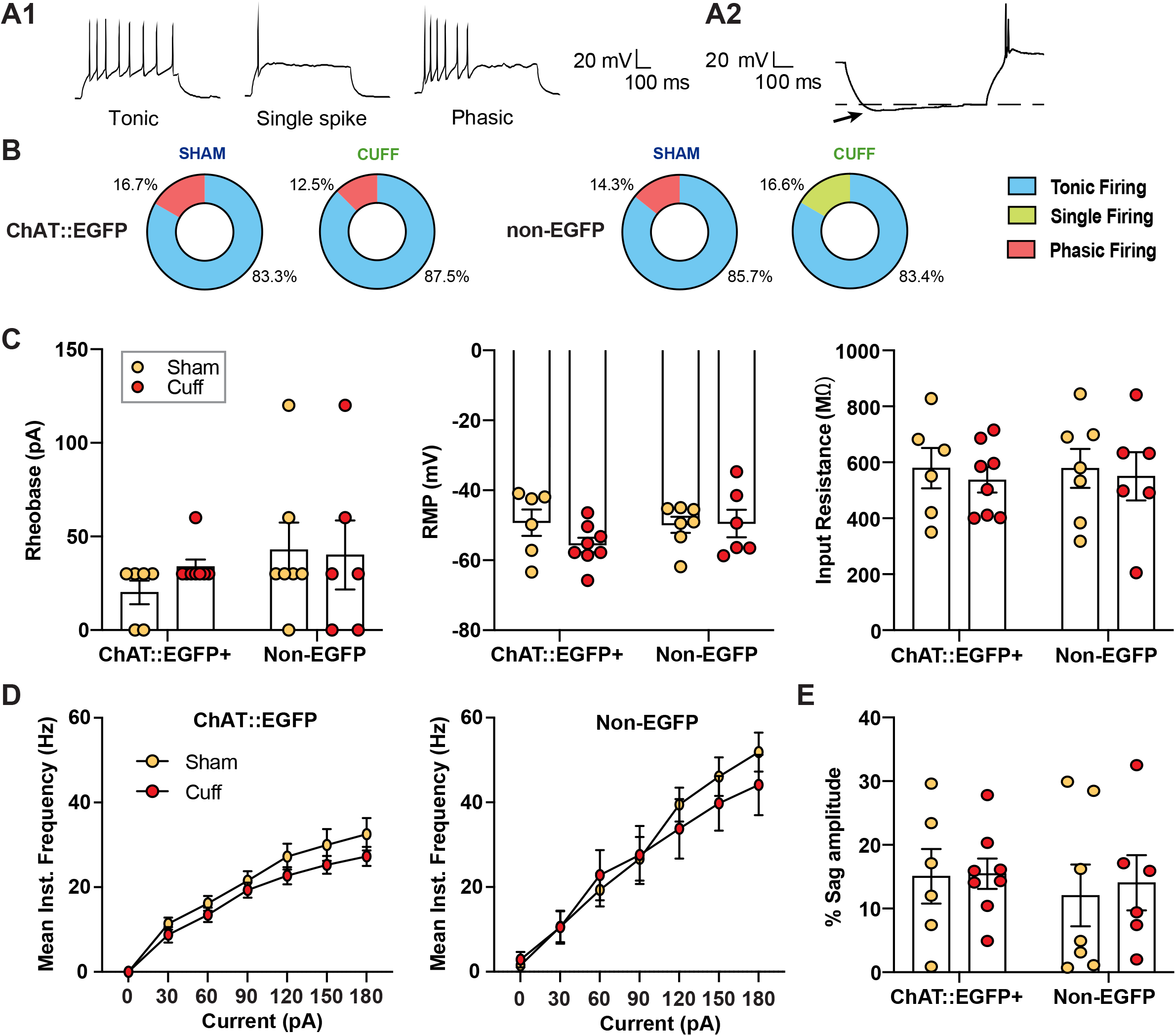
Active and passive properties of lamine III/IV ChAT::EGFP+ and Non-EGFP cells in sham and cuff mice. A: (A1) Recording illustrating 3 different types of firing patterns, observed after injection of 90 pA depolarizing current: tonic, single-spike and phasic, (A2) Example of a response to a hyperpolarizing current showing post-inhibitory rebound. Arrow: hyperpolarizing “sag.” B: The distribution of different firing patterns is not different in the two neuronal populations or as a consequence of the surgery. Surgery-Neuron-Firing pattern interaction: p = 0.6697, 3×3×3 contingency tables. C: Electrophysiological properties of ChAT::EGFP andNon-EGFP cells (Group n = 6-8): Rheobase (3×3×3 contingency tables, p = 0.3765), Resting membrane potential [RMP] (p = 0.2706, Surgery-Neuron interaction: p = 0.253) and Input resistance (p = 0.9191, 2way-ANOVA) were unaltered in both populations following neuropathy. D: Mean Instantaneous frequencies of spikes in function of the injected current, as recorded in ChAT::EGFP and Non-EGFP neurons is similar in sham and cuff animals (Group n = 6-8 neurons). There was a significant effect of neuron type (ChAT::EGFP vs. non-EGFP, p = 1.15e-2, nparLD), current injected (between 0 - 180pA, p = 3.91e-84, nparLD) and a neuron:current injected interaction (comparing different injected currents in ChAT::EGFP and non-EGFP, p = 2.30e-2, nparLD); details in suppl. Fig. 3. There was no Surgery-Neuron-depolarizing current interaction: p = 7.53e-01, nparLD) E: Sag amplitude, expressed as a % of the maximum hyperpolarization of the trace. There was no observable differences in the sag amplitudes (Group n = 6-8, 2way-ANOVA, p = 0.8439).

Altogether, these measures indicate that the network plasticity observed in slices obtained from cuff mice does not alter the excitability of cholinergic neurons.

#### 2.3.3 Number of cholinergic interneurons

There is an on-going debate on whether there is a loss of neurons in the DH that would contribute to the development of hyperalgesia or allodynia following neuropathy. Therefore, we investigated whether DH cholinergic interneurons were affected. We have previously shown that ChAT::EGFP mice are a good model for studying DH cholinergic neurons (Mesnage, et al., 2011). We confirm here that the vast majority (98%) of lamina III-IV EGFP+ are ChAT immunoreactive. We compared the number of EGFP+ neurons in the contralateral and ipsilateral DH of adult ChAT::EGFP+ mice two weeks after surgery. All sections from L3 – L6 spinal segments were taken into account (Fig. 6). On the ipsilateral dorsal horn, there was no statistical difference between the number of ChAT::EGFP+ neurons per 40 μm thick sections in sham (on average 2.13 ± 0.38 neurons) vs. cuff (2.07± 0.38 neurons). This ruled out a change in the number of DH cholinergic neurons after neuropathy.

**Figure 6:**
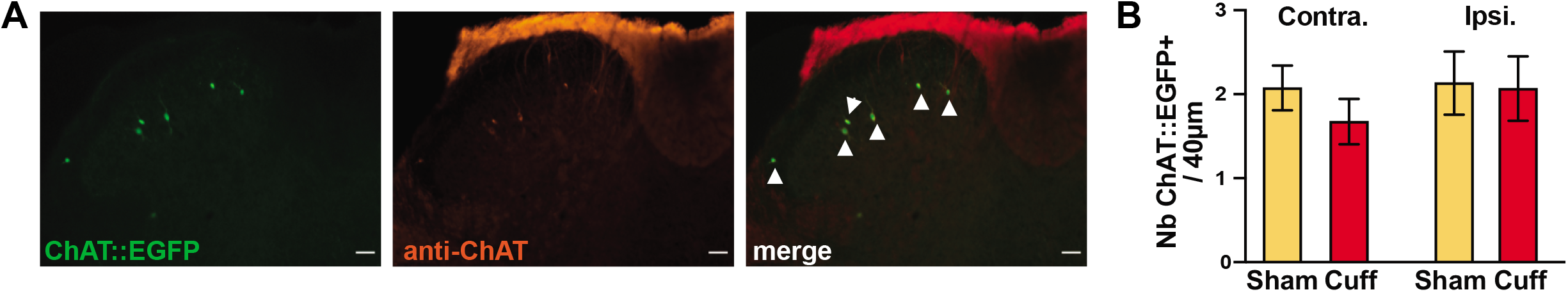
The number of DH cholinergic interneurons is unaltered after cuff surgery. A: Representative illustration of endogeous ChAT::EGFP and anti-ChAT labeling in the ipsilateral dorsal horn of a cuff mouse. The arrowhead denotes overlapping signals (scale = 100 μm). B: Quantification of the total number of DH cholinergic interneurons (ChAT::EGFP fluorescence) in the lumbar (L3-L6) cord of sham and cuff mice [N for Sham and Cuff = 2 mice, 30 sections per condition, 60-64 cells in total per condition]. There is no significant difference between groups (Contralateral side-Surgery-Staining interaction: p = 0.329; 2-way ANOVA | Ipsilateral side-Surgery-Staining interaction: p > 0.99; 2-way ANOVA).

### 2.4 Changes in the downstream network responding to the cholinergic tone

As we found no evidence of specific increased excitation or excitability of DH cholinergic neurons in neuropathic animals, nor a change in neuronal density, we next asked whether the increased impact of the cholinergic tone in these animals could be explained by modifications in direct or indirect targets of cholinergic neurons, i.e. downstream of the tone itself. The cholinergic tone has never yet been described in *in vivo* electrophysiological recordings, nor its consequences on the response of DH neurons to tactile stimulation. We thus first studied these aspects by recording DH neurons *in vivo* in sham and cuff animals to identify a potential substrate to the antinociceptive cholinergic tone and its plasticity, and then turned to pharmacological behavioral experiments.

#### 2.4.1 A neuronal substrate for the antinociceptive cholinergic tone

We performed *in vivo* electrophysiological recordings of lumbar DH neurons in sham and neuropathic animals within the second and third week after surgery. Neurons were selected for their localization in the DH (Fig. 7A) and their response to mechanical stimulation of the hindpaw, either non-nociceptive (touch) and/or nociceptive (pinch) (see methods and (Medrano, et al., 2016)). Local application of 100 μM mecamylamine (on the spinal cord) induced an increased response to mechanical stimulation in neurons recorded from both sham and cuff mice (Fig. 7B). The recorded neurons spanned many DH laminae (Fig. 7A) and the response amplitudes showed high variability, precluding the identification of a potential differential effect of mecamylamine in sham vs. cuff mice. These recordings however demonstrate for the first time *in vivo* the presence of a spinal cholinergic tone, acting through nicotinic receptors, attenuating mechanical responses. This could be a substrate for the antinociceptive spinal cholinergic tone observed in behavioral experiments.

**Figure 7:**
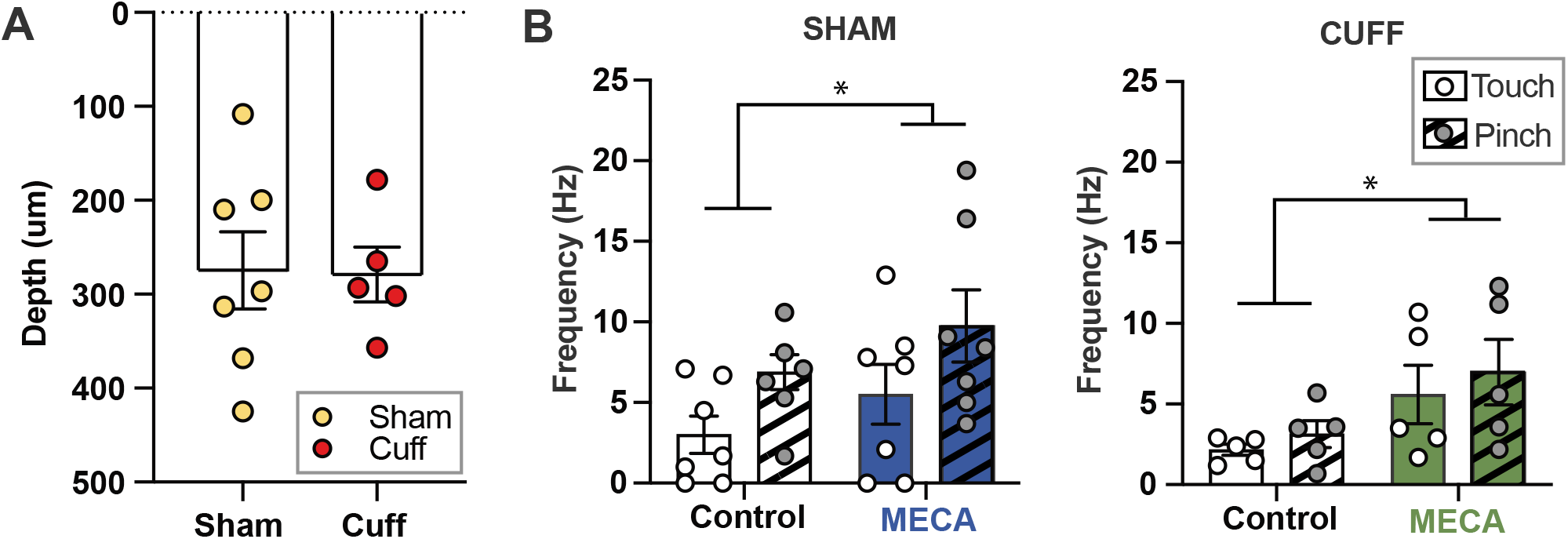
Touch and pinch responses in spinal dorsal horn neurons are modulated by endogenous acetylcholine via nicotinic receptors. A: The depth of recorded DH neurons was similar in sham (274.4 ± 41.06 μm) and cuff (279.0 ± 29.33 μm) mice (sham vs cuff: p = 0.9353; unpaired t-test). B: Mecamylamine increased the intensity of the response of spinal dorsal horn neurons to mechanical stimulation in sham (N = 7 neurons, Drug effect: p = 0.0202; repeated-measures 2 way ANOVA) and cuff mice (N=5 neurons, Drug effect: p = 0.0160; repeated-measures 2 way ANOVA).

#### 2.4.2 Effect of an AChE inhibitor in neuropathic animals

In order to evaluate a potential change of the network downstream of the cholinergic tone, we turned back to behavioral experiments and compared the effect of endogenous spinal ACh in sham and cuff mice by i.t. injection an AChE inhibitor. Physostigmine effect on the contralateral paw was not affected by the surgery (Fig. 8A). For the ipsilateral paw in contrast, the surgery modified the response to physostigmine: in neuropathic mice, 7.5 nmol physostigmine injected i.t. was able to alleviate the mechanical allodynia of the ipsilateral paw while it was ineffective to increase PWT in sham animals (Fig. 8A, at 15 min: 7.5 nmol physostigmine vs. saline p<0.0001 in cuff and p>0.99 in sham; Bonferroni’s multiple comparisons test). The 15 nmol dose however was analgesic for both paws in both sham and cuff mice (Fig. 8A, 15 min: drug vs. saline p<0.0001 for both sham and cuff; Bonferroni’s multiple comparisons test). The dose response curve demonstrates that physostigmine differentially affected the relative mechanical response in the ipsilateral (but not contralateral) paw of cuff vs. sham (Fig. 8B). This favors the hypothesis of a post-synaptic plasticity affecting the downstream players of cholinergic transmission (i.e. post-synaptic receptors).

**Figure 8:**
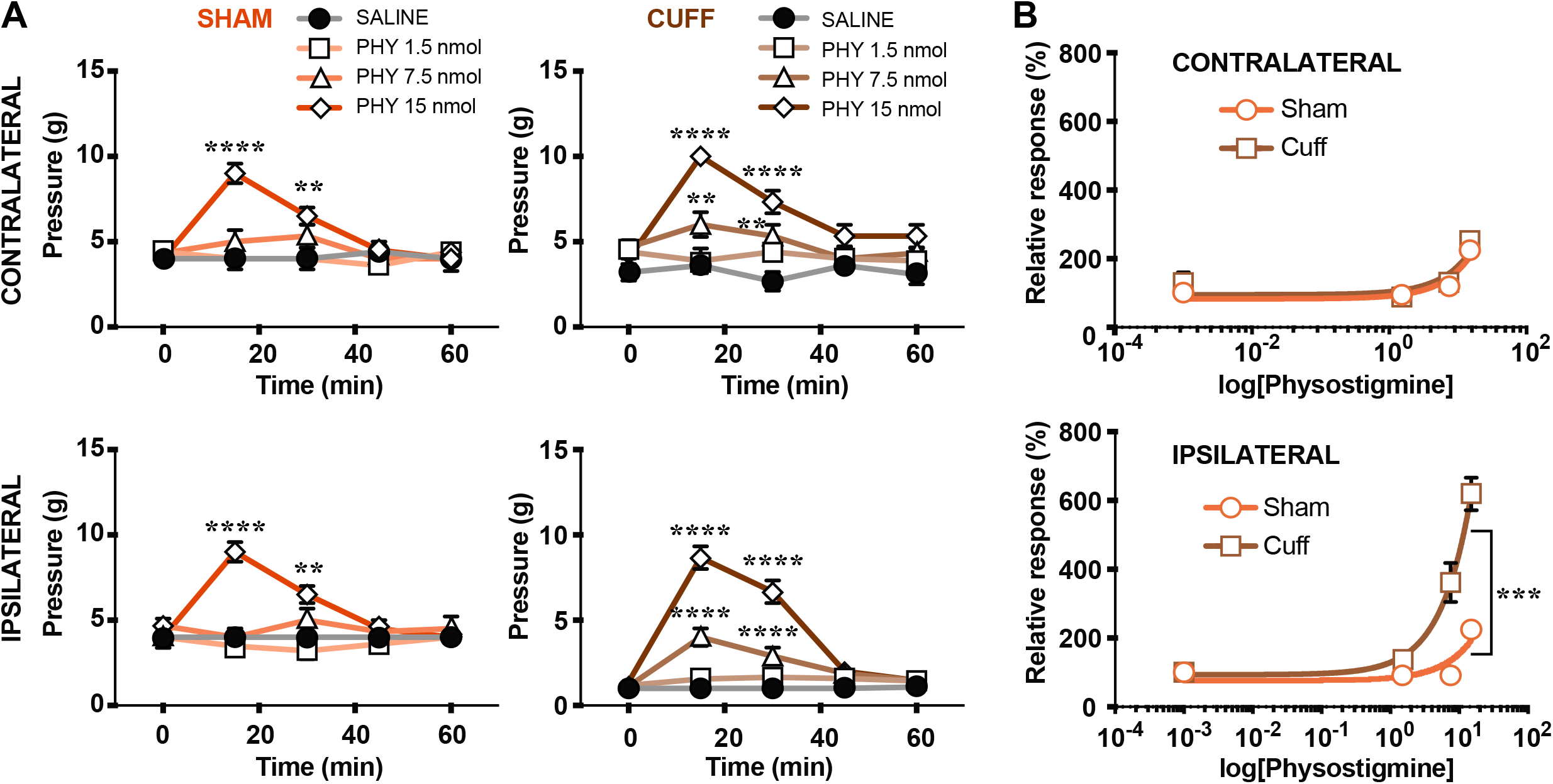
Physostigmine unravels downstream changes to spinal cholinergic tone after neuropathy. A: Time course of the effect of intrathecal physostigmine on the mechanical threshold of sham and cuff mice (Groups n = 3 - 6 mice). (Top) Contralateral PWT was reduced in a dose- and time-dependent manner in sham and cuff mice: significant effect of Drug (p = 9.01e-7) and Time (p = 1.72e-4) but not of Surgery (p = 0.521), and significant interaction between Drug and Time (p = 3.91e-8, nparLD). (Bottom) Idem for the ipsilateral paw: significant effect of Surgery (p = 3.44e-78), Drug (p = 6.26e-22) and Time (p = 5.38e-26), as well as significant interaction between Drug and Time (p = 4.80e-25) and between Surgery, Time and Drug (p = 6.53e-5). The 7,5 nmol physostigmine dose was subthreshold in sham but produced profound anti-allodynia in cuff. **** p < 0.0001, *** p < 0.001 ** p < 0.01, and * p < 0.05, vs. saline, Bonferroni’s multiple comparisons test. B: Dose response curve of physostigmine on PWT of neuropathic animals. The neuropathy induced a shift to the left to the effect of Physostigmine (Ipsilateral side: cuff vs. sham p<0.001; Least square fit)

## 3. Discussion

In this study, we performed an in depth analysis of the antinociceptive spinal cholinergic tone. We demonstrated the persistence and even increased impact of this tone in a model of neuropathic pain, and ruled out changes in lamina III-IV cholinergic neurons as a source of this plasticity. Instead, our data show an increase in the sensitivity of downstream targets to spinal ACh. Low doses of AChE inhibitors, without any effect in control mice, are analgesics in cuff mice which may open new avenues for therapeutical relief of neuropathic pain conditions.

The existence of a “spinal cholinergic tone” has been proposed in the early 1990’s and confirmed since then (Rashid, Furue, Yoshimura, & Ueda, 2006; Rashid & Ueda, 2002; Zhuo & Gebhart, 1991) to describe an endogenous mechanism whose presence is revealed by a pharmacological effect of its antagonization. Although the term is reminiscent of the word “tonic”, it gives no inference on the way ACh is released: either repetitively (through the “tonic firing” of a cholinergic neuron), or phasically (upon a specific stimulation, e.g. nociceptive stimulation (Eisenach, Detweiler, Tong, D’Angelo, & Hood, 1996)). We here similarly define the “cholinergic tone” through the allodynic effect of i.t. mecamylamine injection, suggesting that in basal conditions ACh has an ongoing (or provoked) analgesic effect.

The cuff, a neuropathic pain model chosen for this study, was originally developed for rats (Mosconi & Kruger, 1996) and then adapted for mice (Benbouzid, et al., 2008). In a previous study, we characterized by *in vivo* recordings the loss of inhibitory tone in the spinal DH in this model (Medrano, et al., 2016). We provide here for the first time *in vitro* recordings in the spinal cord of these mice and show that overall, the changes observed in synaptic inputs of DH lamina II neurons in other models are similarly observed in the cuff model. Indeed, we observed a reduction in the frequencies, but not amplitude, of miniature GABA-A currents in lamina II as reported in the spared nerve injury model (SNI) and the chronic constriction injury (CCI) model (Iura, et al., 2016; Moore, et al., 2002). However, we observed no change in the frequency for mEPSCs while a reduction is reported in a study focusing on GABAergic interneurons in the CCI model (Leitner, et al., 2013). Whether this difference is due to a difference in the model, or to the fact that the Leitner et al. study focuses only on GABAergic neurons remains to be established. Importantly, all of our experiments were conducted from the first up to the third week after cuff surgery, a time window where the allodynia phenotype is robustly established but where anxiodepressive-like behaviors are not yet present (Benbouzid, et al., 2008; Yalcin, Bohren, et al., 2011).

A previous study exploring the plasticity of the spinal cholinergic tone used the partial ligation of the sciatic nerve model, and analyzed the thermal paw withdrawal (Rashid & Ueda, 2002). They showed that i.t. injection of 10 nmol mecamylamine had no effect in the injured mice, but higher doses were not tested. In the von Frey mechanical test, we similarly found that 10 nmol mecamylamine was without effect but higher doses exacerbated the allodynia, demonstrating a shift to the right of the dose-response curve of the antagonist. Therefore, rather than a loss of cholinergic tone, our results suggest that its impact is actually increased by the neuropathy.

We also investigated the possible sources of this plasticity. Our previous studies concluded that spinal lamina III-IV cholinergic neurons were the main source of ACh at this level and that their morphology and projections were well suited to impact nociceptive processing, both in rodents and in primates (Mesnage, et al., 2011; Pawlowski, et al., 2013). These cholinergic interneurons are also GABAergic (Mesnage, et al., 2011; Miles, Hartley, Todd, & Brownstone, 2007), and the existence of GABAergic cell death after peripheral neuropathy and its contribution to mechanical allodynia has been proposed although controversially debated (Inquimbert, et al.; Polgar, Gray, Riddell, & Todd, 2004; Scholz, et al., 2005). We took advantage of the ChAT::EGFP mice that have been shown to be a reliable tool to study DH cholinergic neurons (Mesnage, et al., 2011), and found no difference in the number of ChAT::EGFP neurons following neuropathy. Nevertheless, other morphological alterations could exist, as it has been suggested for parvalbumin (PV) interneurons (Petitjean, et al., 2015). Indeed, their number remained unchanged after spared nerve injury in mice, but the number PV+ appositions on PKC-gamma cell bodies was reduced following injury. PV neurons provide feed-forward inhibition onto PKC-gamma neurons thus ‘gating’ the transmission of non-noxious information into nociceptive circuits. This decreased innervation has therefore been proposed to be a substrate of mechanical allodynia (Petitjean, et al., 2015).

Beyond changes in density or morphology, a modification of the synaptic inputs received by the cholinergic interneurons could explain a change in the spinal cholinergic tone. A combination of recent reports demonstrate that Aβ inputs gain access to LI projection neurons through the loss of multiple feedforward inhibitory circuits following nerve injury (Abraira, et al., 2017; Alba-Delgado, et al., 2015; Cheng, et al., 2017; Foster, et al., 2015; Imlach, et al., 2016; Lu, et al., 2013; Peirs, et al., 2015; Petitjean, et al., 2015). Although the network organization of the dorsal horn starts to be elucidated (Cordero-Erausquin, Inquimbert, Schlichter, & Hugel, 2016), the position of cholinergic neurons with respect to this feedforward inhibitory circuit is still unknown. In the cuff model, we observed a decrease in inhibition in lamina II neurons. Those lamina II neurons could be inhibitory neurons projecting to deeper laminae (Santos, Rebelo, Derkach, & Safronov, 2007), and their disinhibition could thus explain the increased frequency of inhibitory currents observed in laminae IIIIV. This increased inhibition of cholinergic interneurons does not translate into decreased excitability, indicating that the release of ACh might not be significantly impaired. Overall, the membrane properties of these neurons were unchanged in neuropathic animals. Similar observations are reported for LIII (GAD67 or not) or LII GABAergic neurons (Gassner, Leitner, Gruber-Schoffnegger, Forsthuber, & Sandkuhler, 2013; Schoffnegger, Heinke, Sommer, & Sandkuhler, 2006), suggesting that changes in membrane excitability or altered firing patterns in the spinal cord dorsal horn are unlikely causes for the alterations underlying neuropathic pain or for the observed plasticity of the cholinergic tone.

As the analysis of DH cholinergic interneurons did not explain the increased impact of the cholinergic tone in neuropathic animals, we then explored downstream mechanisms, in the spinal nociceptive network. Previous *in vivo* recordings performed in rats demonstrated that projection neurons in the cervical cord had an increased response to mechanical stimulation after topical application of an AChE inhibitor (neostigmine) (Chen & Pan, 2004). Using similar recordings in mice, but on unidentified DH neurons, our study illustrates that a nicotinic antagonist has the opposite effect on mechanical responses, demonstrating for the first time a potential substrate the analgesic cholinergic tone *in vivo*. Our recordings also show that this tone is still present after cuff injury, but no differential effect of the cholinergic tone after neuropathy is revealed.

To finally identify the plasticity that could explain the increased impact of the cholinergic tone in cuff mice, we analyzed the analgesic properties of increasing i.t. doses of an AChE inhibitor, physostigmine, with the von Frey test. These experiments demonstrated a shift to the left of the dose response curve of this drug since doses that were subthreshold in sham mice returned the PWT in the cuff to pre-surgery levels. This is reminiscent of data obtained in the rat after systemic (i.p.) injection of another AChE antagonist, donepezil, in the spared nerve injury model (Kimura, Hayashida, Eisenach, Saito, & Obata, 2013) demonstrating that 0.6 and 1 mg/kg of i.p. donepezil were anti-hyperalgesic in a mechanical paw withdrawal test, while they showed no effect before the ligature. Concerning the thermal sensory modality, i.t. injections of AChE inhibitors are also analgesic in naïve rats (Chiari, et al., 1999; Naguib & Yaksh, 1994), and a study performed in mice showed again the anti-hyperalgesic effect of very low doses of i.t. neostigmine (another AChE inhibitor, 3ng) in the tail flick test in the SNL model, while higher doses (from 115 ng i.t.) were required for analgesia in control mice (Takasu, et al., 2006). Interestingly, nicotinic agonists also have analgesic properties after i.t. injection in the PNL mouse model, both in mechanical and thermal tests, at doses (10 nmol nicotine, 0.3 nmol epibatidine) inefficient in sham mice (Rashid & Ueda, 2002). This, together with our results, strongly suggests that major changes occur at the receptor level, either in density or in composition.

Alterations in the expression of nAChRs have been reported in neuropathic conditions. The expression of the α6 nAChR subunit in murine DRGs was demonstrated to correlate with the mechanical allodynia produced after neuropathy (Wieskopf, et al., 2015), while the mRNA level of the α5 and β2 nAChR subunits is increased in neuropathic rats (Yang, et al., 2004). A specific study of alterations of nAChRs on the targets of cholinergic neurons in the dorsal horn of the spinal cord should help elucidate the changes occurring in the mouse cuff model and possibly explaining the plasticity of the cholinergic tone. In this study, we have focused on the spinal nicotinic tone because we have revealed the cholinergic tone through antagonization of nAChRs by mecamylamine. Mecamylamine is a general nicotinic antagonist (Papke, Sanberg, & Shytle, 2001), but i.t. hexamethonium or the α4β2 nAChR antagonist DHβE, or the α3β2*/α6β2* nAChR antagonist, α-CTX-MII, have been shown to have a similar effect in mice or rats (Rashid, et al., 2006; Rashid & Ueda, 2002; Yalcin, Charlet, et al., 2011; Young, Wittenauer, McIntosh, & Vincler, 2008). In addition, the allodynic effect of nicotinic antagonists is lost in nAChR β2 knock-out mice (this study and (Yalcin, Charlet, et al., 2011)), further confirming the involvement of β2-containing nAChRs. In contrast, the α7 nAChR antagonist MLA (10 nmol) has no effect, suggesting that α7 nAChRs are not involved in the cholinergic tone in mice (Rashid, et al., 2006; Rashid & Ueda, 2002; Yalcin, Charlet, et al., 2011). However, it should be noted that the increased ACh level induced by AChE inhibition could act both on nicotinic and muscarinic post-synaptic receptors. In naïve rats, the analgesic effect of neostigmine on thermal responses has been shown to be mediated by muscarinic receptors in males but muscarinic and nicotinic receptors in females (Chiari, et al., 1999; P. M. Lavand’homme & Eisenach, 1999). However muscarinic antagonists (atropine, pirenzepine, 4-DAMP, or methoctramine) had no effect on von Frey responses when injected i.t. in naïve mice (Honda, et al., 2002; Paqueron, Conklin, & Eisenach, 2003), suggesting that there might be species or sensory modality differences.

In this study, we have demonstrated that the spinal cholinergic tone modulating nociceptive responses was still present, and even more efficient, after peripheral nerve injury. Our data do not support a change in the number or electrophysiological properties of dorsal horn cholinergic interneurons, but rather favor a modification occurring at the post-synaptic site. Overall, the increased analgesic effect of i.t. AChE antagonists in neuropathic conditions is promising. Until now, AChE inhibitors have mainly been used in the clinics in situations of acute pain (parturition or post-operative (Eisenach, 2009)). The fact that lower doses (thus with fewer side effects) could be efficient in chronic pain conditions opens new avenues for the treatment of neuropathic pain.

## 4. Material and methods

### 4.1 Animals

This study utilized wild-type and transgenic CD1 and C57BL/6 mice between 3 – 10 weeks. Two different transgenic lines have been used: ChAT::EGFP CD1 (von Engelhardt, Eliava, Meyer, Rozov, & Monyer, 2007) and β2*-nAChR knock-out (KO) C57BL/6J mice (Picciotto, et al., 1995). The animals were group-housed between two to six animals per cage and maintained on a 12-hour light/dark cycles with food and water provided *ad libitum*. The animal facility Chronobiotron UMS3415 is registered for animal experimentation under the Animal House Agreement B6748225. All protocols were approved by the “Comité d’Ethique en Matière d’Expérimentation Animale de Strasbourg” (CREMEAS, CEEA35) and the Ministère de l’enseignement supérieur, de la recherche et de l’innovation under the reference 02729.01.

### 4.2 Cuff surgery and behavioral assessment

The cuff model is used to induce neuropathic pain in mice (Benbouzid, et al., 2008). The surgery was carried out under ketamine (1 μl/g-Imalgene1000, Merial), azepromazin (0,60 μl/g - Calmivet, Vetoquinol) and medetomidine (1,18 μl/g - Domitor, Orion pharma/Elanco) anesthesia. The surgery was performed as previously described (Yalcin, et al., 2014). Briefly, a 2 mm section of split PE-20 polyethylene tubing (Harvard apparatus, Les Ulis, France) was placed around the right common branch of sciatic nerve in the cuff mice group. Sham-operated mice underwent the same surgical procedure without cuff implantation. After performing the surgery, Atipamezol chlorhydrate (Antisedan, Orion pharma/Vetoquinol) mix (10 μl/g of animal), an antidote to medetomidine, was provided to speed up the recovery from the anesthesia. After a one-week recovery period, the neuropathic phenotype was assessed with von Frey tests.

Intrathecal (i.t.) injections were performed under gaseous anesthesia (3% Isoflurane for induction and 2% for maintenance) as previously described (Yalcin, Charlet, et al., 2011). We assessed the role of spinal cholinergic modulation on mechanical transmission in naïve and neuropathic CD1 mice with von Frey filaments (Yalcin, et al., 2010). Mice were habituated for 15 minutes in clear Plexiglas boxes (7cm x 9 cm x 7 cm) on an elevated mesh screen. Filaments of increasing diameter were slowly brought to the plantar surface of each hind paw and removed after a slight bent. The mechanical threshold is determined as the first filament inducing at least three withdrawals out of five consecutive trials. The results were expressed in grams. For naïve animals, the left and right paw data were averaged for each time point. The mice were tested prior to injection or surgery to establish the baselines. A minimum of three tests were performed over 3 to 7 days to determine the mechanical withdrawal threshold. Motor coordination was tested using a rotarod (Bioseb). Mice were trained on 2 consecutive days where they ran at constant speed of 8 and 16 rotations per minute (rpm) without falling for at least 60 secs. On the experimental day, the rod was programmed to accelerate from 0 to 40 rpm over 300 s. Each mouse was allowed three trials where the average rpm and time at the point of failure was recorded to establish the baseline. Subsequently, mecamylamine was intrathecally injected and the mice were tested again 3 times starting after 15 min post-injection.

### 4.3 In vitro electrophysiological recordings

From 7 to 24 days post-surgery, ChAT::EGFP CD1 mice were anesthetized with a mix of i.p. ketamine (200mg/kg - Imalgene 1000, Merial) and xylazine (20 mg/kg - Rompun 2%, Bayer). The animals were transcardially perfused for approximately 3 minutes with ice-cold sucrose-based Artificial Cerebral Spinal Fluid (ACSF) composed of (in mM): 252 Sucrose, 2.5 KCl, 2 CaCl2, 2 MgCl2, 10 Glucose, 26 NaHCO3, 1.25 NaH2PO4, 2 kynurenic acid, continuously bubbled with carbogen (95% O_2_/ 5% CO_2_). Two approaches were used to extract the spinal cord: hydraulic extrusion (Chery & De Koninck, 1999) or laminectomy (Flynn, Brichta, Galea, Callister, & Graham, 2011). A 200 to 300 μm horizontal slice (HS) was performed, containing the first four laminae of the DH. We also performed 350 to 400 μm thick transverse slices (TS). The slices were allowed a one-hour recovery period prior to recording at room temperature in oxygenated ACSF containing (in mM): 126 NaCl, 2.5 KCl, 2 CaCl2, 2 MgCl2, 10 Glucose, 26 NaHCO3, 1.25 NaH2PO4.

Cholinergic neurons were identified by the presence of the enhanced green fluorescent protein (EGFP) produced under the control of the cholineacetyltransferase (ChAT) promoter. Cells were recorded in the lumbar cord, in the area corresponding to termination of the sciatic nerve (end of L3 – L6 spinal segments). In HS, LIII/IV was identified by few ChAT::EGFP neurons located on the surface of the first slice (positioned upside-down). In TS, LIII/IV was identified as the laminae just ventral to the translucent substantia gelatinosa (LII), or ventral to the EGFP+ cholinergic plexus (Mesnage, et al., 2011). For voltage clamp experiments, a cesium (Cs) based intracellular solution was used (in mM): 80 Cs2SO4, 5 KCl, 2 MgCl2, 10 HEPES, 10 Biocytin (pH = 7.3). The cells were maintained at −60 mV and 0 mV for excitatory and inhibitory post-synaptic currents respectively. For current clamp recordings, a K-based intracellular solution was used (in mM): 136 CH3KO3S, 2 MgCl2, 10 HEPES, 10 Biocytin. For both intracellular solutions, the liquid junction potential was not corrected. The recordings were made through the Axopatch 200A (Axon instruments) or the Multiclamp 700A (Molecular devices). The current traces were filtered at 5 kHz before digitalization via a digitizer (BNC-2110, National Instruments) at 20 kHz. Data acquisition was performed using WinEDR and WinWCP softwares (Strathclyde Electrophysiology Software, John Dempster, University of Strathclyde, Glasgow, UK).

For voltage clamp, the synaptic events (recorded over a 5-minute period) were detected with the threshold search method with WinEDR software. The threshold was set, for miniature and spontaneous EPSCs, to absolute amplitudes ≥ 2 pA and a duration ≥ 1.25 ms. For miniature and spontaneous IPSCs, the thresholds were set to absolute amplitudes ≥ 3 pA and a duration ≥ 1.75 ms. A baseline track-time of 5 ms was implemented prior to each event. Following the automatic detection by the software, all captured events were inspected individually, and only events that possessed a quick rise followed by an exponential decay were kept. The absolute frequency is calculated as the number of events over the total duration of the recording. The peak amplitudes were measured for accepted events via semi-automated procedures in WinWCP. However, individual events were excluded if they contained overlapping events or had an unstable baseline. The frequency and amplitude of currents were analyzed over a 5-minute duration. For current clamp, the firing patterns were classified as previously reported (Prescott & De Koninck, 2002).

### 4.4 In vivo electrophysiological recordings

Adult CD1 males were anesthetized with urethane (2.5 g/ kg, i.p.) and the trachea was cannulated. Rectal temperature was continuously monitored, and the animal was maintained at 35.5°C using a heating pad (TC-1000; Bioseb). The mouse was placed in a stereotaxic and spinal frame with 2 clamps fixed on its vertebra to immobilize the vertebral column. A laminectomy was performed to expose L3-L5 segments of the spinal cord, and a small chamber (approximately 0.1 ml) was created with 2% agar around the exposed lumbar spinal cord. After removal of the dura, the spinal cord was covered with saline (NaCl 0.9%) or with drugs diluted in saline (see below).

Single-unit extracellular recordings from spinal dorsal horn neurons were performed as previously described (Medrano, et al., 2016). Only neurons responding to mechanical stimulation of the ipsilateral hind paw were included in this study. Recordings were made with a glass electrode (Harvard Apparatus, Holliston, MA) filled with 0.5M CH3COOK (resistance: 15-25 mV). A motorized micromanipulator (Narishige, Tokyo, Japan) was used to gradually descend the electrode with 4-mm steps until the single-unit activity of a neuron was recorded. The recording electrode was inserted at depths ≤500 mm from the surface of the spinal cord (corresponding to lamina I to V). The signal was amplified (IR-183; Cygnus Technology, Delaware Water Gap, PA), filtered at 0.3 to 3 kHz (Brownlee amplifier; AutoMate Scientific, Berkeley, CA), and digitized at 20 kHz with a MICRO3-1401 (CED, Cambridge, United Kingdom). Data were analyzed offline with Spike 2 software (CED). The cutaneous receptive field of the recorded neuron was identified by touching the ipsilateral hind paw. The response to a non-nociceptive mechanical stimulus, touch, and to a nociceptive one, pinch, was determined as described before (Medrano, et al., 2016)). The touch stimulus was applied by brushing the skin with a camel’s hair brush for 10 seconds (8-10 times). The pinch stimulus was applied by means of small serrated forceps (Graefe forceps; Fine Scientific Tools, Vancouver, Canada) for 10 seconds.

Mecamylamine hydrochloride (nicotinic receptor antagonist; Sigma (St. Louis, MO, USA)) was applied directly at the surface of the dorsal horn using the agar chamber described above. Stock solutions were first prepared in water at 100x their final concentration, stored at −20 °C and, on the day of the experiment, diluted in saline to their final concentration. The drug was applied topically onto the recording site of the spinal cord after careful removal of saline of the pool. Ten minutes following drug application, the response of neurons to the mechanical stimuli was tested (Medrano, et al., 2016). All solutions were applied at 34-35°C.

### 4.5 Tissue fixing and staining

The animals were transcardially perfused under pentobarbital (Ceva Sante Animal, 54.7mg/ml, i.p. injection) with 0.1M phosphate buffer (PB, pH 7.4) followed by 4% Paraformaldehyde (PFA) in PB for 15 minutes. The lumbar part of the spinal cord was sliced in 40μm-thick transverse sections using a vibrating blade microtome (VT 1000S, Leica, Rueil-Malmaison, France), and serially collected in wells. The sections were washed three times with PBS and saturated with PBS/ 0.5% Triton X-100/ 5% donkey serum at room temperature for 45 minutes. The sections were incubated at room temperature overnight in PBS/ 0.5% Triton X-100/ 1% donkey serum with a goat polyclonal ChAT antibody (1:500 dilution, Chemicon Millipore, AB 144P). After three PBS washes, the slides were incubated with a CY3 anti-goat antibody (1:400 dilution, Jackson Immunoresearch) for 2 hr. The sections were finally washed three times with PBS and mounted with fluorescent mounting media (Dako, Les Ulis, France). Slides were examined under fluorescence using a microscope (Leica) and an exhaustive counting of fluorescent neurons was made in both the contralateral and ipsilateral DH in all sections between L4 and L6.

### 4.6 Statistics

The statistical analysis was performed with the statistical software R (version 3.4.1) in conjunction with Graphpad software (Prism 7 for Mac, GraphPad Software, Inc., San Diego, CA, USA). For *in vitro* data, almost all statistical comparisons for frequency, amplitude and cell properties, considering the various dependent variables, were made with the ANOVA function. The original or transformed (log or inverse function) datasets were verified to show a normal distribution (Shapiro-Wilk normality test). For post-hoc comparisons, interactions with only one or two variables were made with the Bonferroni test with multi-comparison correction. A more complex 3×3×3 contingency table was performed online at http://vassarstats.net (Log-Linear Analysis for an AxBxC Contingency Table found under *Frequency data* tab) for firing pattern, rebound spike observations and rheobase. For *in vivo* and *in vitro* data passing requirements for parametric tests, statistical evaluation was carried out with repeated measures 2-way ANOVA. In addition, the depths for *in vivo* recordings were compared with unpaired t-test.

As the vonFrey data is a discrete ordinal data, it did not pass the conditions to use parametric tests. We thus analyzed it using the non-parametric test “nparLD”, provided as a plug-in by the statistical software R (Noguchi, Gel, Brunner, & Konietschke, 2012). nparLD provides an ANOVA type multiple-factor analysis taking into account a longitudinal variable (time) along with multiple dependent (right vs left paw) and independent (surgery; drug treatment) variables. The Bonferroni test was also used for post-hoc multi-comparisons against saline.

The significance level was set at P <0.05 and data were expressed as mean ± SEM for graphs.

## Acknowledgements

We gratefully acknowledge the support from the University of Strasbourg Institute for Advanced Study (USIAS) and ANR-13-JSV4-0003-01 GRANT to MCE. DD is the recipient of a Region Alsace doctoral fellowship, YML of a Fondation pour la Recherche Médicale post-doctoral fellowship, and SK was supported by the NeuroTime Erasmus Mundus Joint Doctorate Neuroscience PhD program (funded by the European Commission), and CB a recipient of a fellowship from the French Ministry of Higher Education and Research. The authors express their gratitude towards J.L. Rodeau for his expert assistance on the statistical approaches, Philippe Isope and Yves De Koninck for critical readings; and Sophie Reibel-Foisset and the Chronobiotron (UMS 3415, Centre National de la Recherche Scientifique) for mice handling. The authors have no conflicts of interest to declare.

