## Supplementary figures for "Enhanced analgesic cholinergic tone after neuropathy"

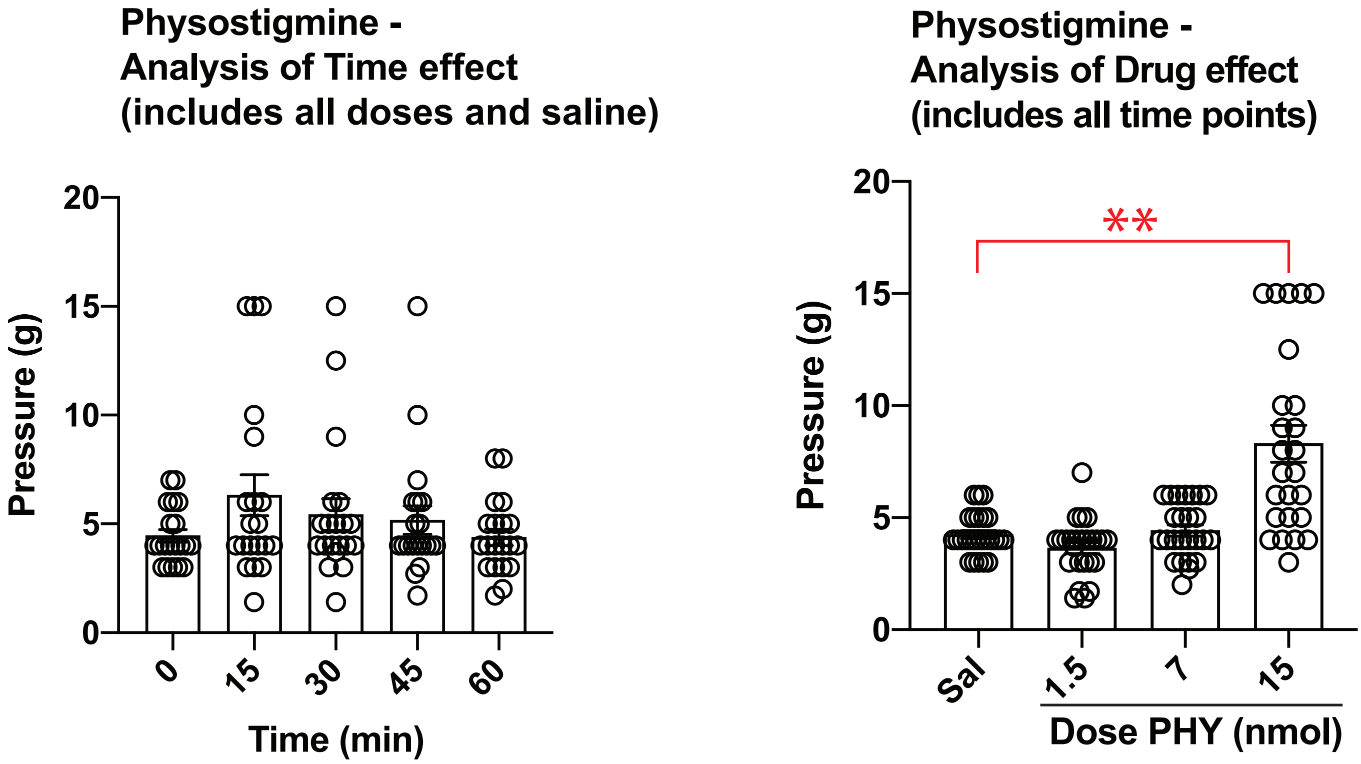


**Suppl. Fig. 1:**
**Statistical analysis of the effect of i.t. Physostigmine on the Von Frey test.**
Shown are the analysis of the two factors: Time (left) and Drug (right). There was no statistical effect of Time, but an effect of Drug (p=2.43e-3, nparLD).
** $p < 0.01$ Bonferroni's multiple comparisons test.

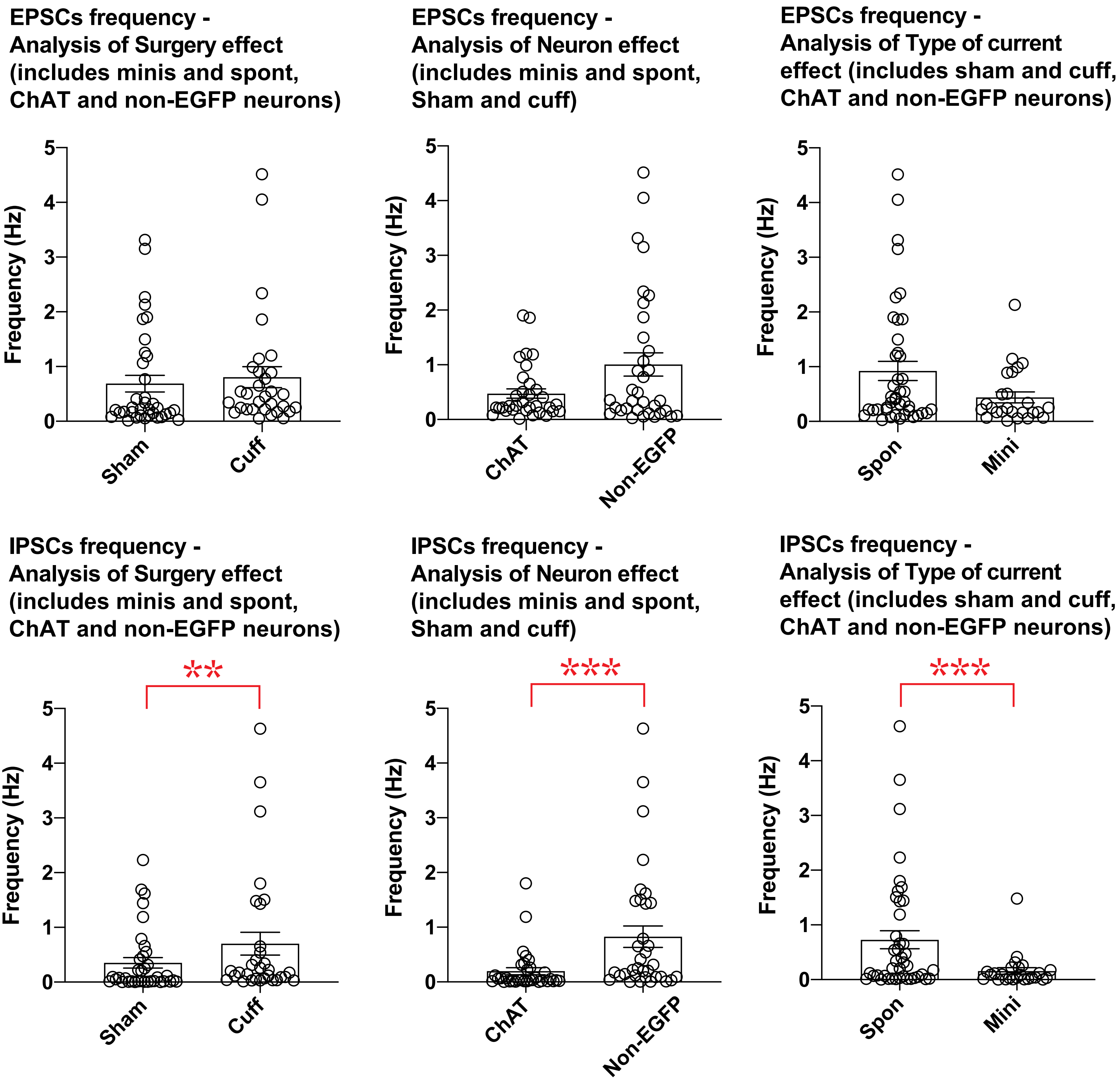


**Suppl. Fig. 2:**
**Statistical analysis of the frequency of synaptic currents in DH lamina III–IV neurons (horizontal slices, sham and cuff mice)**
Shown are the analysis of the three factors: surgery (left), type of neuron (ChAT::EGFP or non-ChAT, middle) and type of current (spontaneous vs. miniature, right), for EPSCs (top) and IPSCs (bottom).
There was no statistical difference in EPSCs frequencies between groups.
For IPSCs, main effects were significant: neuron (ChAT::EGFP vs. non-EGFP, $p = 0.000179$, 2-way ANOVA), surgery (sham vs. cuff, $p = 0.004616$, 2-way ANOVA) and type of current (spontaneous vs. miniatures, $p = 1.051e-3$, 2-way ANOVA).

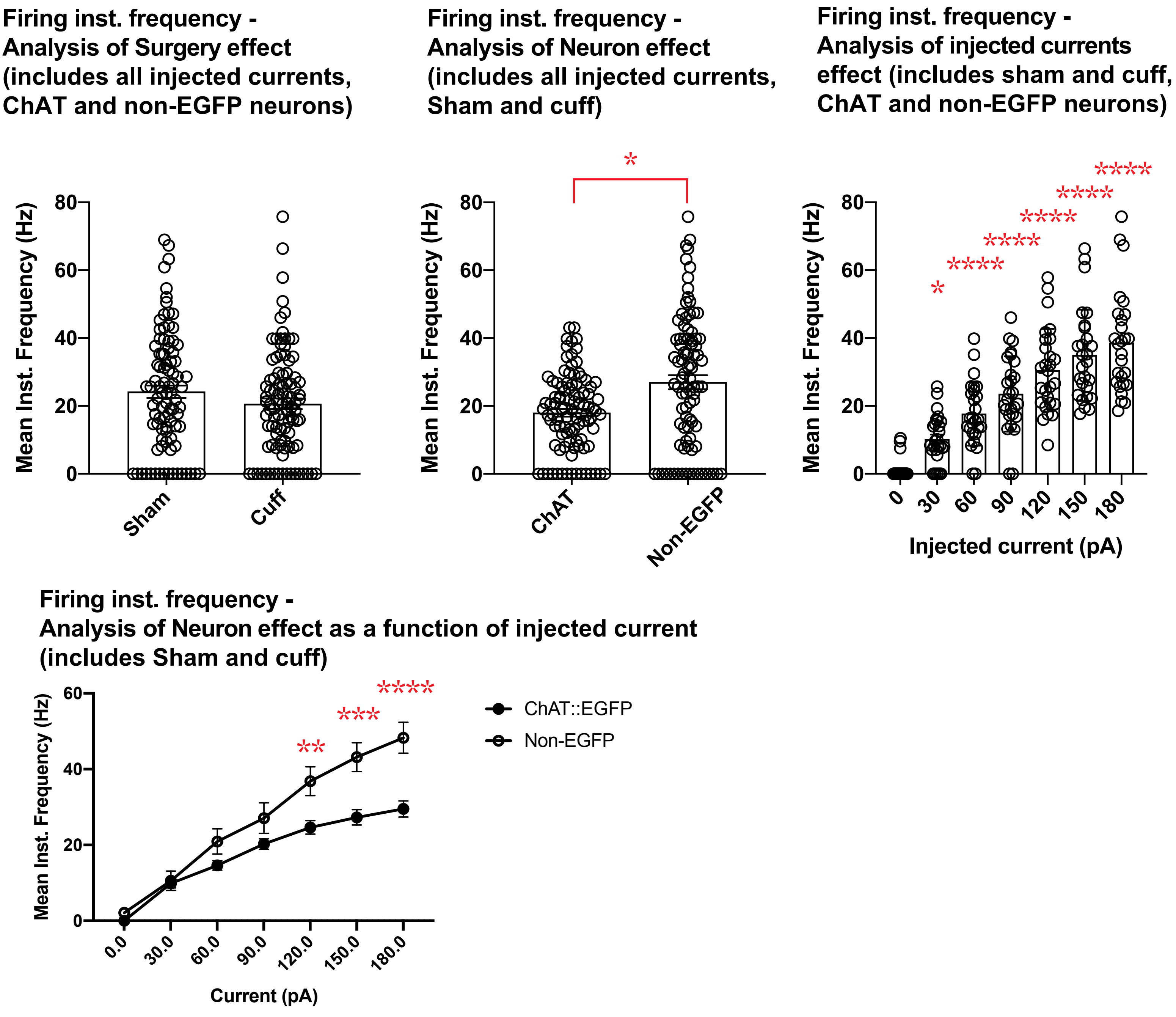


**Suppl. Fig. 3:**

**Statistical analysis of the mean instantaneous frequencies of spikes induced by current injections (horizontal slices, sham and cuff mice)**

**Top:** Shown are the analysis of the three factors: surgery (left), type of neuron (ChAT::EGFP or non-ChAT, middle) and injected current (0–180pA, right). Main effects were significant: neuron (ChAT::EGFP vs. non-EGFP, p = 1.15e-2, nparLD), current injected (between 0–180pA, p = 3.91e-84, nparLD). Post-hoc analysis for current injection vs. 0 pA: * p < 0.05, *** p < 0.0001. Not illustrated in the graph, comparisons vs. 30 pA: p < 0.001 for 90 pA, and p < 0.0001 for 120 to 180 pA; comparisons vs. 60 pA: p < 0.001 for 120 pA, and p < 0.0001 for 150–180pA; comparisons vs. 90 pA: p < 0.01 for 150 pA, and p < 0.0001 for 180 pA)

**Bottom:** There was an interaction between the factors Neuron and Current injected (p = 2.30e-2, nparLD). **** p < 0.0001, *** p < 0.001 ** p < 0.01 vs. ChAT::EGFP neurons, Bonferroni's multiple comparisons test.
